# Dynamin-2 promotes Atg9A retrieval from phagophores during autophagy

**DOI:** 10.64898/2026.03.11.711183

**Authors:** Andrew Caliri, Alejandro Martorell Riera, Akash Saha, Panagiota Kolitsida, Cinta Iriondo Martinez, Samuel Itskanov, Janos Steffen, Carla M. Koehler, Alexander M. van der Bliek

**Affiliations:** Department of Biological Chemistry, David Geffen School of Medicine at UCLA; Department of Chemistry and Biochemistry, UCLA

## Abstract

Autophagy involves the rapid growth of phagophores through membrane addition. This growth is triggered by vesicles containing the Atg9A protein. However, Atg9A is not incorporated into mature autophagosomes. We now demonstrate that Dynamin-2 (Dnm2) colocalizes with the BAR domain protein Endophilin-B1 (EndoB1/Bif-1/SH3GLB1) and other autophagy proteins when autophagy is induced. Our data suggest that Atg9A is retrieved from phagophores via fission, with Dnm2 acting as the membrane scission protein. Blocking Atg9A recycling, either by mutating Dnm2, using RNA interference, or applying chemical inhibitors, results in Atg9A remaining in autophagosomes and being degraded during autophagy. Overall, these findings provide new insights into the roles of membrane-scission proteins in autophagy.

## Introduction

Autophagy is a universal process for removing damaged components in the cell and recycling proteins during starvation (Choi AM et al, 2013). Much has been learned about this process through studying various protein complexes that quickly and intricately assemble autophagosomes, which then fuse with lysosomes for degradation (Hurley JH & Young LN, 2017, Levine B & Klionsky DJ, 2017, Yamamoto H et al, 2023). Autophagosomes are double-membrane compartments that capture cargo destined for lysosomal breakdown (Bento CF et al, 2016). In the early stages of autophagy, precursors called phagophores form near the endoplasmic reticulum (Karanasios E et al, 2016, Orsi A et al, 2012). These phagophores rapidly grow to enclose the material to be degraded in autolysosomes (Chang C et al, 2021, Noda NN, 2021).

The triggers for autophagy are either starvation or ubiquitinated cargo, such as defective mitochondria and other organelles marked for degradation by autolysosomes. Autophagy receptors like p62 and NDP52 connect ubiquitinated cargo to FIP200 and ULK1 (Ravenhill BJ et al, 2019, Turco E et al, 2019), beginning the process that leads to autophagosome formation. Autophagosome membranes originate from Atg9A-containing vesicles coming from the Golgi, recycling endosomes, and other organelles (Sawa-Makarska J et al, 2020). These Atg9A vesicles interact with FIP200 and ULK1 during early stages of autophagy (Stanley RE et al, 2014). They serve as seeds for autophagosome growth (Olivas TJ et al, 2023). The vesicles expand into a larger envelope around the cargo by adding membrane (Gomez-Sanchez R et al, 2018). Atg2 acts as a conveyor, extracting lipids from the ER via interactions with VMP1 and TMEM41B (Yamamoto H et al, 2023), then transferring these lipids to the outer leaflet of the phagophore membrane (Maeda S et al, 2019, Osawa T et al, 2019, Valverde DP et al, 2019). To balance the growth of the outer and inner leaflets, excess lipids on the outer surface are moved to the inner surface by Atg9A, which functions as a scramblase (Guardia CM et al, 2020, Maeda S et al, 2020, Matoba K et al, 2020).

Atg9A is also the only known autophagy protein with transmembrane segments. Interestingly, this protein was found not to be incorporated into mature autophagosomes (Orsi A et al, 2012). Instead, Atg9A is likely retrieved from growing phagophores, but the mechanism remains unknown.

In this study, we examined the potential roles of Dynamin-2 (Dnm2) in autophagy. Dnm2 is one of three classic dynamins in mammalian cells (Antonny B et al, 2016, Ferguson SM & De Camilli P, 2012, Schmid SL & Frolov VA, 2011). It is expressed throughout the body and facilitates membrane scission in various endocytic processes, including the formation of both clathrin- and caveolin-coated vesicles (Conner SD & Schmid SL, 2003). Dynamin-1 and dynamin-3 are mainly found in neurons, where they function at pre- and post-synaptic densities during endocytosis (Ferguson SM & De Camilli P, 2012). Over time, several additional roles for Dnm2 have been suggested. These include a role in creating exocytic vesicles that bud from the TGN (Jones SM et al, 1998, Liu YW et al, 2008), forming actin comets on endocytic vesicles (Lee E & De Camilli P, 2002, Orth JD et al, 2002), aiding a late stage of cytokinesis (Thompson HM et al, 2002), supporting a late stage of mitochondrial fission (Lee JE et al, 2016), regenerating autolysosomes during lipid droplet breakdown in hepatocytes (Schulze RJ et al, 2013) and forming ATG9A-containing vesicles from recycling endosomes during autophagy (Soreng K et al, 2018, Takahashi Y et al, 2016). Many of these proposed functions still require further validation, but Dnm2’s role in endocytosis has been observed across diverse contexts.

All three dynamins have very similar sequences. Each contains five distinct protein domains: an N-terminal GTPase domain, a middle domain, a pleckstrin homology (PH) domain, a GTPase effector domain (GED), and a proline-rich domain (PRD) (Ferguson SM & De Camilli P, 2012). The middle domain and GED fold back on each other to form a stalk, while the PH domain interacts with membranes, and the PRD binds to SH3 domains in various adaptor proteins (Chappie JS et al, 2010, Faelber K et al, 2011, Ford MG et al, 2011, Gao S et al, 2010). A large pool of dynamin proteins exists as homomeric tetramers in the cytosol, but a small fraction gathers at specific spots on the plasma membrane to promote membrane scission (Schmid SL & Frolov VA, 2011). Multiple contacts between adjacent dynamin molecules at those sites enable assembly into chains, which then form multimeric spirals that wrap around the necks of budding endocytic vesicles and constrict to sever the membrane while hydrolyzing GTP (Schmid SL & Frolov VA, 2011).

One key factor in guiding dynamins to different membranes is whether specific binding partners are present. Dynamins interact with PI(4,5)P_2_ at the plasma membrane (Achiriloaie M et al, 1999) and with SH3 domain-containing proteins like amphiphysin and endophilin A during endocytosis (Simpson F et al, 1999). Cooperative binding between dynamin and amphiphysin or endophilin A has been shown to help tubulate membranes *in vitro* and facilitate membrane severing *in vivo* (Farsad K et al, 2001, Meinecke M et al, 2013, Renard HF et al, 2015). Both amphiphysin and endophilin A have isoforms, such as amphiphysin-2 (Bin-1) and endophilin B1 (EndoB1, SH3GLB1 or Bif1), which might act on other membranes (Farsad K et al, 2001, Lee E et al, 2002). EndoB1 has been linked to autophagy, either as part of an autophagy protein complex (Takahashi Y et al, 2007) or by affecting Atg9A vesicle formation at recycling endosomes (Takahashi Y et al, 2016). While exploring EndoB1’s role in autophagy, we observed colocalization of EndoB1 and Dnm2, prompting us to further investigate Dnm2’s potential role in autophagy. In this study, we show that Dnm2 facilitates the retrieval of Atg9A from phagophores.

## Results

### Dnm2 and EndoB1 colocalize during autophagy

To test for Dnm2-EndoB1 colocalization, we performed immunofluorescence experiments on wild-type MEF cells treated with or without rapamycin to induce autophagy. These cells were permeabilized with digitonin to enhance the signal from membrane-bound proteins by removing soluble cytosolic proteins (Liu SH et al, 1998). We observed a significant increase in Dnm2 and EndoB1 colocalization after rapamycin treatment (Fig 1A), as shown by Mander’s correlation coefficient (Fig 1B) and scatter plots (Figs S1A and B). Similar results were obtained using Pearson’s correlation coefficient (not shown). To further evaluate these interactions, we performed live cell imaging of transfected proteins with fluorescent tags, as demonstrated with images (Fig 1C), Mander’s coefficient (Fig 1D), and scatter plots (Figs S1C and D). In each case, rapamycin and CCCP significantly increased colocalization between Dnm2 and EndoB1.

**Figure 1.**
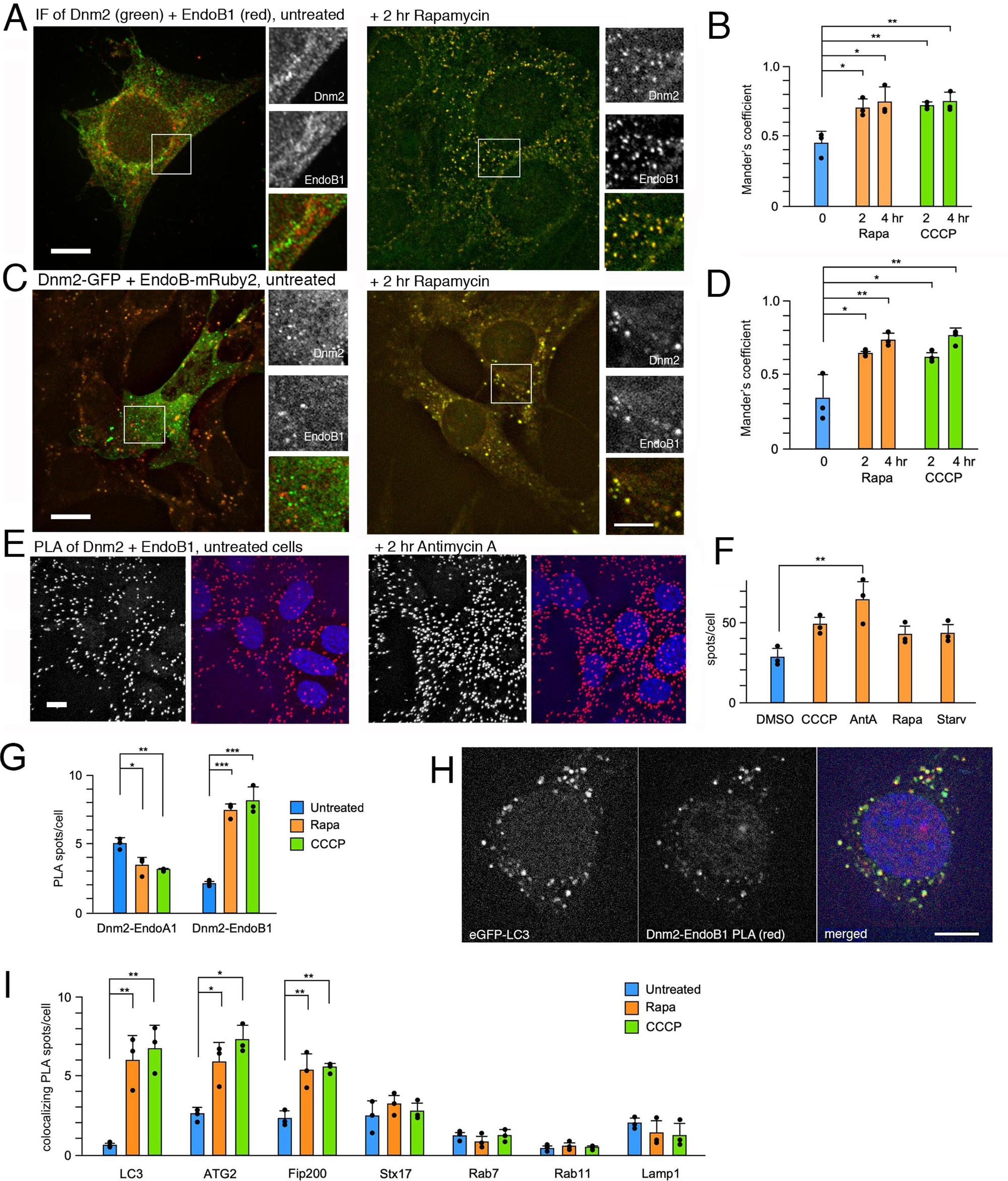
Colocalization of Dnm2 and EndoB1 with autophagy proteins. **(A)** Immunofluorescence of endogenous Dnm2 (green) and EndoB1 (red) in MEFs is shown without or with 2 hr rapamycin. To reduce the diffuse cytosolic staining of Dnm2, these cells were incubated with 0.05% digitonin before fixation with paraformaldehyde. Scale bar is 10 μm for whole cells and 5 μm for enlarged portions. **(B)** Mander’s coefficients for colocalization shown for untreated cells and after 2 or 4 hr with rapamycin or CCCP. Average and SD are shown for three biological replicates, each determined with 5 technical repeats. Statistical analysis used one-way ANOVA with Tukey’s post hoc comparisons of the biological replicates. **(C,D)** Similar experiments with transfected cells expressing Dnm2-GFP and EndoB1-mRuby2. **(E)** Colocalization of Dnm2 and EndoB1 tested with Proximity Ligation Assay (PLA) shown with or without 2 hr Antimycin A treatment. Scale bar is 10 μm. **(F)** Average numbers of PLA dots/cell shown for Dnm2 and EndoB1 colocalization in untreated MEFs, MEFs treated with CCCP, Antimycin A, rapamycin, and starvation to induce autophagy. Numbers of dots were counted in 50 cells/experiment. Average and SD are shown for three biological replicates, each determined with 10 technical repeats. Statistical analysis used one-way ANOVA with Tukey’s post hoc comparisons of the biological replicates. **(G)** Relative numbers Dnm2-EndoB1 PLA spots increase, while Dnm2-EndoA1 PLA spots marginally decrease when autophagy is induced. Average and SD are shown for three biological replicates, each determined with 25 technical repeats. Statistical analysis used one-way ANOVA with Tukey’s post hoc comparisons of the biological replicates. **(H)** Images of PLA spots for endogenous Dnm2 and EndoB1 (red) and colocalization with EGFP-LC3 (green) when mitophagy is induced (4 hr CCCP). Scale bar is 10 μm. **(I)** A larger fraction of Dnm2-EndoB1 PLA spots colocalize with GFP-tagged LC3, Atg2 and Fip200 when autophagy is induced with rapamycin or CCCP, suggesting that Dnm2 is connected with autophagy. The fraction that colocalizes with GFP-tagged Lamp1 decreases and remains low for Stx17, Rab11 and Rab7. Average and SD are shown for three biological replicates, each determined with 13-25 technical repeats. Statistical analysis used one-way ANOVA with Tukey’s post hoc comparisons of the biological replicates.

We further examined the colocalization of endogenous Dnm2 and EndoB1 proteins using a proximity ligation assay (PLA). First, we confirmed the specificity of PLA with MEF cells harboring homozygous deletions of the Dnm2 and EndoB1 genes, as well as MEF cells lacking both genes. Western blots verified the absence of Dnm2 and EndoB1 expression in these cell lines (Fig S1E). The PLA for Dnm2 and EndoB1 in these knockout cells did not produce any spots (Figs S1F), indicating that the PLA with Dnm2 and EndoB1 antibodies is highly specific. Additionally, no PLA spots were observed when using antibodies against Fis1, a mitochondrial outer membrane protein facing the cytosol, or Hsp60, a mitochondrial matrix protein, suggesting that the distance between the cytosol and mitochondrial matrix is sufficient to prevent PLA signal (Fig S1G). In contrast, PLA with antibodies for the confirmed partners MFF and Drp1 (Otera H et al, 2010) yielded numerous spots, confirming the effectiveness of this technique (Fig S1G).

PLA was then used to assess the colocalization of Dnm2 and EndoB1 under various autophagy-inducing conditions. We observed significant increases in PLA spots with each autophagy-inducing condition (Figs 1E and F), which matched the colocalization seen through immunofluorescence and with fluorescent-tagged proteins. The increase in PLA spots under autophagy-inducing conditions, however, was consistent with the increased colocalization observed in immunofluorescence of endogenous and transfected proteins. We therefore conclude that the colocalization of Dnm2 and EndoB1 increases during autophagy.

### Colocalization of Dnm2 and EndoB1 with phagophores

Although the number of Dnm2-EndoB1 PLA spots increased when autophagy was induced, the number of Dnm2-EndoA1 PLA spots decreased, indicating a shift from endocytosis to autophagy (Figs 1G). To further support a potential role in autophagy, we assessed colocalization with other autophagy proteins using PLA and fluorescent proteins. We observed a significant increase in colocalization of Dnm2-EndoB1 with EGFP-LC3 upon induction of autophagy (Figs 1H and I). We then examined colocalization of PLA spots with several other autophagy-related proteins. Results, summarized in Fig 1I, show additional increases in the fraction of Dnm2-EndoB1 PLA spots associated with two early autophagy markers (Atg2 and Fip200). No changes were seen in colocalization with Stx17 or Rab7, which are found on mature autophagosomes. There was a decrease in the fraction of PLA spots associated with LAMP1, which may indicate a shift away from the role in autophagolysosomal recycling previously observed (Schulze RJ et al, 2013). We conclude that a large portion of Dnm2-EndoB1 PLA spots are located at or near sites of phagophore formation, but this association diminishes during later stages of autophagy.

### No effects of Dnm2 deletion on mitophagy or starvation-induced macro-autophagy

Since Dnm2 and EndoB1 colocalize during autophagy induction, we tested whether these proteins contribute to the process. We first examined the effects of Dnm2 deletion on CCCP-induced mitophagy, as previous studies have linked Dnm2 to mitochondria (Karbowski M et al, 2004, Lee JE et al, 2016). Super-resolution structured illumination microscopy (SIM) images of EGFP-LC3 and mitochondrial DsRed show cup-shaped phagophores surrounding mitochondria in Dnm2 KO cells after 6 hours of CCCP treatment (Fig 2A). These phagophores look the same as in wild-type cells. Fluorescence images reveal CCCP-induced mitochondrial fragmentation in all cells, along with gradual mitochondrial removal over 24 hours in both WT and Dnm2 (Fig 2B). A similar pattern is observed in Western blot analysis of the mitochondrial matrix protein Hsp60 in Dnm2 KO cells (Figs 2C and D), indicating that CCCP-driven Hsp60 degradation occurs over 24 hours. If anything, degradation seems faster in Dnm2 KO.

**Figure 2.**
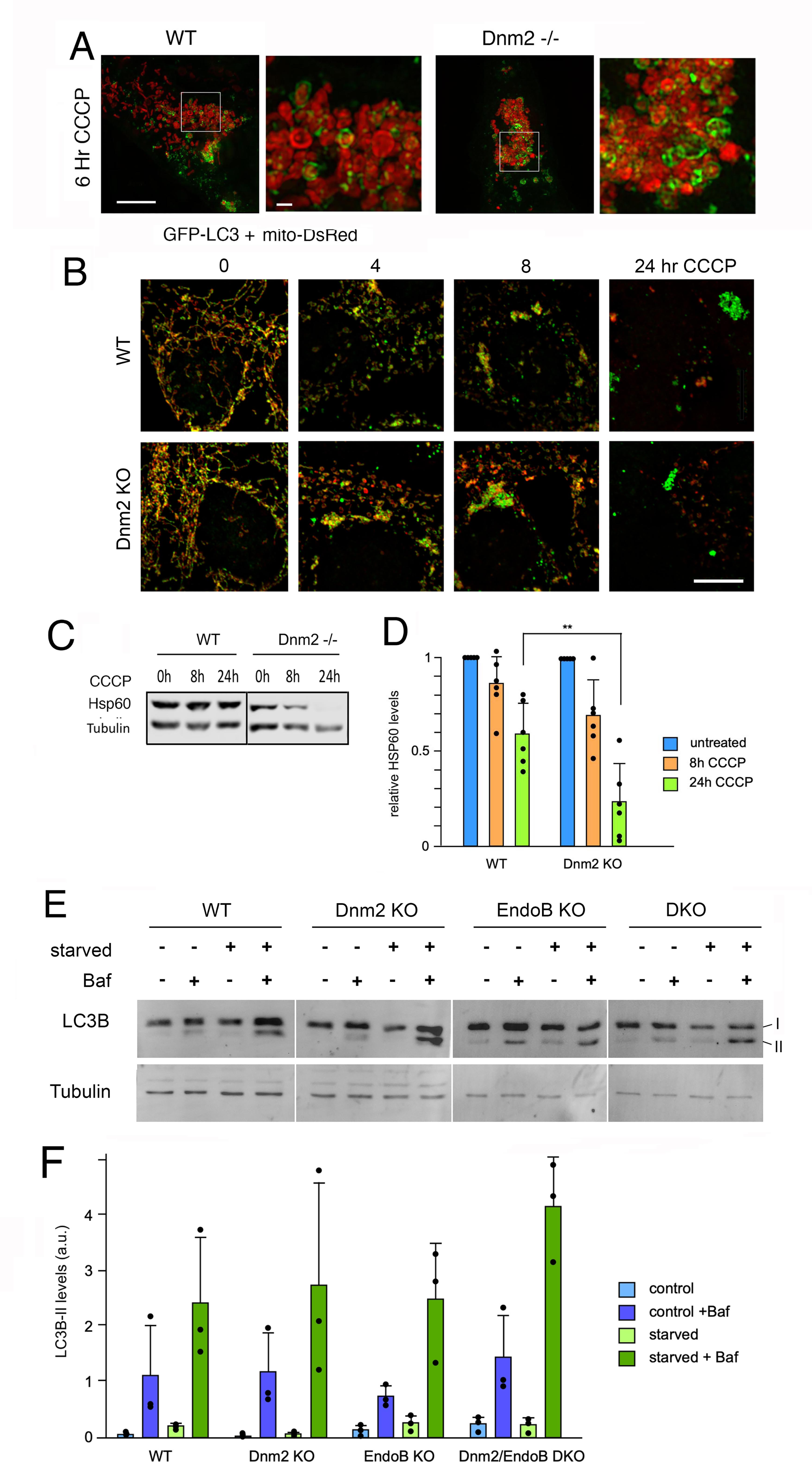
Little effect of Dnm2 and EndoB1 deletions on mitophagy and macro-autophagy. **(A)** Super-resolution images (SIM) show the formation of LC3-membranes (green) encapsulating mitochondria (red) in WT and Dnm2 KO MEFs treated with CCCP. Cells were treated for 6 hr with 20 μM CCCP. Mitochondria were detected with mitoDsRed and autophagic membranes were detected with EGFP-LC3. Scale bar is 10 μm for the overview and 1 μm for the enlargements. **(B)** Effects of CCCP on mitophagy in WT and Dnm2 KO MEFs observed with immunofluorescence microscopy of fixed cells using Tom20 (red) and Hsp60 (green) antibodies. At 24 hr, most mitochondria are degraded. Scale bar is 10 μm. **(C)** Western blots showing the effects of CCCP on Hsp60 levels in WT and Dnm2 KO MEFs. **(D)** Average intensities of Hsp60 bands in four independent experiments, relative to tubulin levels and normalized to the 0-hr time point. Average and SD are shown for six biological replicates. Statistical analysis used one-way ANOVA with Tukey’s post hoc comparisons. **(E)** Western blots showing the effects of starvation-induced macroautophagy and Bafilomycin A treatments on LC3B lipidation in wildtype, Dnm2, EndoB KO and DKO MEFs. **(F)** Average intensities of LC3B-II bands in four independent experiments, relative to tubulin levels, expressed in arbitrary units (a.u.). Average and SD are shown for three biological replicates. Statistical analysis used one-way ANOVA with Tukey’s post hoc comparisons.

To evaluate whether deletions of Dnm2 or EndoB affect macroautophagy, cells were starved for four hours in EBSS, then processed for cell extraction and Western blot analysis of LC3B lipidation. The blots show no significant differences in LC3B-II levels among Dnm2, EndoB KO, or DKO MEFs (Figs 2 E and F). Similar experiments with Dnm2 and EndoB KO HeLa cells also found no differences (Figs S2 A and B). We conclude that mitophagy and macroautophagy proceed normally in Dnm2 and EndoB KO cells.

### Transfer of LC3 to autophagolysosomes is unimpeded in Dnm2 KO cells

To monitor downstream events affected by mutations in Dnm2 during autophagy, we transfected cells with RFP-GFP-LC3, which appears as yellow spots in the cytosol and red spots when autophagosomes fuse with lysosomes, as lysosomal acidification quenches GFP. Our results show that Dnm2 and EndoB KO cells exhibit more red fluorescence spots than wild-type cells after CCCP treatment (Figs 3A and B). Autophagosomes in Dnm2 and EndoB KO cells move more quickly toward fusion with lysosomes compared to those in wild-type cells treated with CCCP (Fig 3C). To further investigate effects on autophagy, we measured the colocalization of LC3 and Rab7 after autophagy was triggered by Rapamycin. Colocalization of LC3 and Rab7 was not reduced in Dnm2 KO cells (Figs 3D and E), indicating that autophagosome formation occurs normally in these cells.

**Figure 3.**
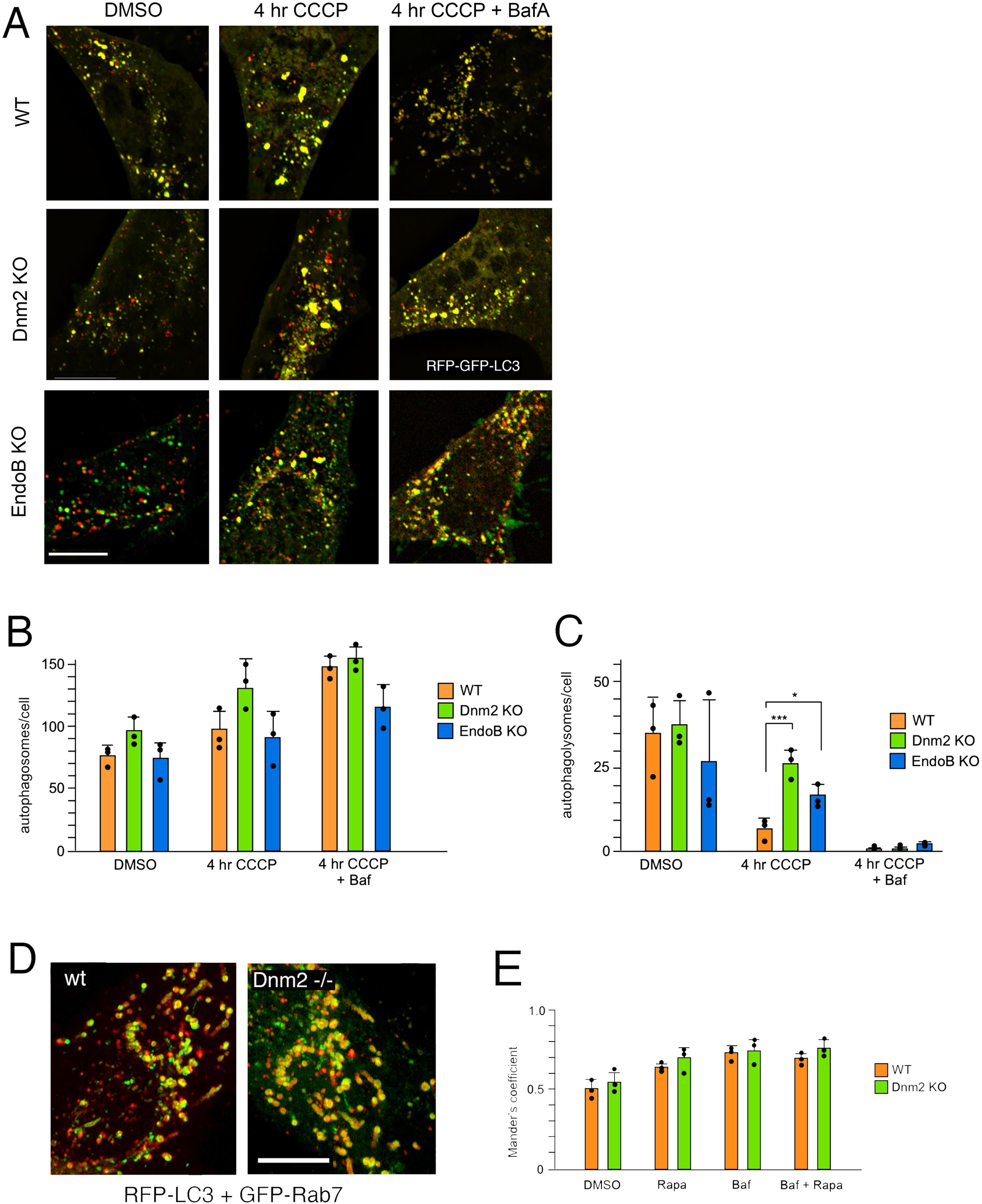
Transfer of LC3 to lysosomes is unimpeded. **(A)** Tracking autophagy with RFP-GFP-LC3 marker in WT and in Dnm2 and EndoB KO MEFs shows transfer of LC3 to autophagolysomes, visible as red spots due to acidic quenching of GFP. Scale bar is 10 μm. **(B)** Numbers of phagophores or autophagosomes per cell, detected as yellow spots, in WT and in Dnm2 and EndoB KO cells. Average and SD are shown for three biological replicates, each determined with 5 technical repeats. Statistical analysis used one-way ANOVA with Tukey’s post hoc comparisons, but no significance was detected. **(C)** Numbers of autophagolysosomes, detected as red spots, are significantly increased in Dnm2 and EndoB KO MEFs treated with CCCP. Statistics as in panel B. **(D)** Live cell imaging of the effects of wildtype and Dnm2 KO cells showing colocalization of RFP-LC3 and GFP-Rab7. In this case, autophagy was induced for 1 hr with Rapamycin. Scale bar is 10 μm. (**E**) Levels of LC3 and Rab7 colocalization expressed as Mander’s coefficient for untreated cells and cells treated with Rapamycin, Bafilomycin, or both. Average and SD are shown for three biological replicates, with 7 technical replicates per biological replicate. Statistical analysis used one-way ANOVA with Tukey’s post hoc comparisons, but no significance was detected.

### Atg9A is retained on LC3-containing membranes in Dnm2 KO cells

The Atg9A protein, which helps deliver membrane to growing phagophores, is usually not incorporated into mature autophagosomes, suggesting a recycling process retrieves this protein. We studied the potential effects of Dnm2 KO on the retrieval of Atg9A from autophagic membranes, since this process involves membrane scission at phagophores. We checked whether the retrieval blockage could be detected using fluorescent proteins. Dnm2 KO and wild-type cells were transfected with RFP-tagged Atg9A and EGFP-LC3 and then fixed for microscopy. We observed clear colocalization of Atg9A-myc and HA-LC3 by IF in fixed Dnm2 KO cells, but not in wild-type cells (Fig 4A), confirmed by Mander’s coefficients (Fig 4B). Similar changes were RFP-Atg9A and GFP-LC3 in live cells (Figs S3A and B). There was also a decrease in spots labeled only with Atg9A, not LC3 (Fig S3C), supporting the idea that fusion with phagophores reduces the supply of Atg9A vesicles. Changes in Atg9A and LC3 colocalization were also observed with the dynamin inhibitor dynasore (Figs S3B and C). The changes were also enhanced by treatment with Bafilomycin A (Figs S3D and E), in WT cells transfected with Dnm2 siRNA (Figs S3F-H), and in Dnm2 KO HeLa cells (Figs S3I-K), confirming that Dnm2 is necessary to prevent Atg9A and LC3 from colocalizing in autophagosomes.

**Figure 4.**
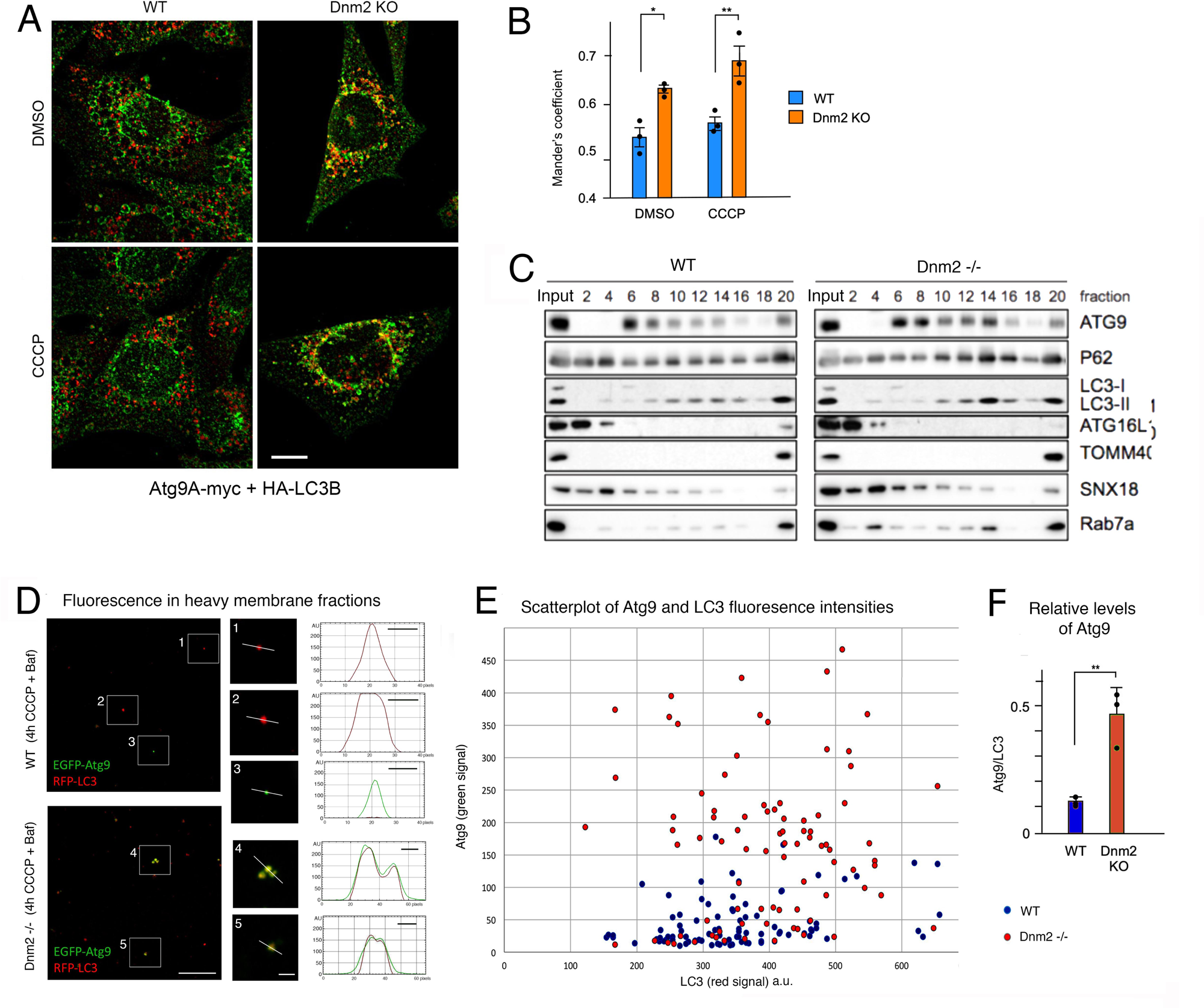
Atg9A remains associated with LC3 in Dnm2 KO cells. **(A)** Fixed cell fluorescence images showing stably expressed Atg9A-myc colocalizes with HA-LC3 in Dnm2 KO MEFs treated with CCCP, but not in wild type MEFs. Scale bar is 10 μm. **(B)** Histogram showing Mander’s coefficient for colocalization of Atg9A-myc and HA-LC3 in WT and Dnm2 KO MEFs after incubation with DMSO or CCCP. Average and SD are shown for three biological replicates, with five technical replicates per biological replicate. Statistical analysis used one-way ANOVA with Tukey’s post hoc comparisons. **(C)** Subcellular fractionation of homogenates from WT and Dnm KO MEF cells, which were treated for 4 hr with Rapamycin. Membranous organelles were separated with a 1-22% Ficoll density gradient and analyzed with Western blots. The bottom fraction (#20) contains mitochondria and other dense organelles such as mitophagosomes, autophagolysosomes or lysosomes. Atg9A is normally present in light fractions (fraction #6), but partially shifts to an intermediate density (fraction #14) in Dnm2 KO cells, along with some p62, LC3-II and Rab7a. **(D)** Images of autophagosome-containing fractions from WT and Dnm2 KO cells transfected with EGFP-Atg9A and RFP-LC3, followed by 4hr treatments with CCCP and Bafilomycin to induce mitophagy and to accumulate autophagosomes. Autophagosomes were enriched by differential centrifugation. Enlargements show individual spots with line tracings to plot fluorescence intensities. Scale bar is 10 μm in the overview images and 2 μm in the enlargements. The x-axis in the fluorescence intensity plots show numbers of pixels (24 pixel/μm) and the bars show 500 nm. **(E)** Scatter plot of peak fluorescence intensities for individual spots, as shown in panel F (arbitrary units). **(F)** Histogram of the peak fluorescence intensities of Atg9A in LC3 spots relative to the peak intensities of Atg9A in spots without LC3. Average and SD are shown for three biological replicates with 30 technical replicates each. Statistical analysis used one-way ANOVA with Tukey’s post hoc comparisons.

Next, we used subcellular fractionation to detect potential abnormal Atg9A-containing organelles in Dnm2 KO cells. The distribution of these organelles was analyzed using Ficoll density gradients, as described previously (Orsi A et al, 2012). Autophagy was induced in WT and Dnm2 KO MEF cells with rapamycin instead of CCCP because the mitophagosomes formed by CCCP could not be separated from intact mitochondria in Ficoll gradients. In contrast, rapamycin promotes the formation of less dense autophagosome-related structures that can be separated with Ficoll gradients. Atg9A is usually found in a light membrane fraction (fraction #6 in Fig 4C), but some protein is detected in the heavy membrane fractions of induced cells (fraction #20). Other autophagy proteins, including p62 and LC3-II, are present in this fraction alongside mitochondrial (TOMM40) and endo/lysosomal (Rab7a) proteins. Importantly, Dnm2 KO cells show an additional buildup of certain proteins (Atg9A, p62, LC3-II, and Rab7a) in a new fraction (fraction #14 in Fig 4C). This fraction does not contain Atg16L1, which acts early to promote LC3 lipidation, nor does it include SNX18, which was previously suggested to work with Dnm2 in Atg9A vesicle formation at early endosomes (Soreng K et al, 2018). These findings suggest that fraction #14 contains abnormal autophagosomes with Atg9A, LC3-II, Rab7a, and p62. Notably, Atg9A in fraction #14 is significant because this protein is normally absent at a stage when Atg16L1 is removed and autophagosomes fuse with Rab7a-containing compartments.

To further investigate whether Atg9A and LC3 co-exist in the same compartments in Dnm2 KO cells, we imaged individual autophagosomes from crude biochemical isolates as described (Gao et al., 2010b).

MEF cells were transfected with EGFP-Atg9A and RFP-LC3, then treated for 4 hours with CCCP and Bafilomycin to induce mitophagy and promote autophagosome accumulation. These autophagosomes were enriched through differential centrifugation and washed twice in 1.0 M NaCl to dissociate vesicle clusters held together by ionic interactions. Fractions of these samples were mounted on slides and examined using fluorescence microscopy. Extracts from WT cells showed fluorescent spots containing LC3 but little or no Atg9A, likely representing mature autophagosomes (Fig 4D). Occasionally, spots with Atg9A but lacking LC3 were observed, which may be Atg9A vesicles or organelles. In contrast, extracts from Dnm2 KO cells consistently displayed spots with overlapping LC3 and Atg9A fluorescence (Fig 4D). Selected line tracings for these spots are shown in Fig 4D, with a third example for Dnm2 KO cells shown in Figs S4A-C. The LC3 and Atg9A fluorescence intensity tracings closely matched in Dnm2 KO extracts, indicating that membranes containing both proteins had fully merged. A scatterplot of peak intensities for various spots showed that many in Dnm2 KO extracts contain both proteins, whereas this was not the case in WT extracts (Fig 4E).

Additionally, a histogram of peak intensities from three independent experiments revealed significant differences in their distribution in Dnm2 KO cells (Fig 4F).

We hypothesized that a Dnm2-dependent process could retrieve SNARE proteins along with Atg9A vesicles, and that this retrieval would be impaired in Dnm2 KO cells. We examined the localization of VAMP7, a v-SNARE that is usually found in late endosomes but is also associated with Atg9A vesicles (Aoyagi K et al, 2018, Yu L et al, 2017). Our results show increased colocalization of VAMP7 with LC3 in Dnm2 KO cells, and this colocalization increases upon CCCP-induced mitophagy (Figs S4D and E), suggesting that VAMP7 is also retrieved from phagophores via Dnm2-mediated membrane scission.

### Dnm2 promotes the retrieval of Atg9A from autophagosomes

We used live-cell imaging to explore whether Dnm2 localizes to sites of Atg9A retrieval from autophagosomes. We tracked RFP-tagged Dnm2 near or on phagophores labeled with BFP-LC3 and vesicles labeled with GFP-Atg9A. Even in untreated cells, we observed RFP-Dnm2 spots positioned between BFP-LC3-labeled phagophores and GFP-Atg9A–labeled vesicles. These Dnm2 spots disappeared within a second, and the organelles separated. Examples are shown in Fig 5A. The frequency of these events was low (1 in 3 videos for untreated cells), likely due to limitations of the imaging system (50 images of a cell segment in a single plane at 350-msec intervals). However, extrapolating to entire cells suggests about 25 events per cell per minute, consistent with a role in autophagy. Due to photobleaching and rapid organelle movement, we could not determine if multiple fission events happen on growing phagophores. Fission at the plasma membrane can be blocked by the dynamin inhibitor Dynole 34-2 and promoted by the dynamin activator Ryngo 1-23 (Jackson J et al, 2015, Lasic E et al, 2017). We tested how these chemicals affected the appearance and disappearance of RFP-Dnm2 spots between BFP-LC3- and GFP-Atg9A-labeled organelles. We observed a significant decrease in Dnm2 spots with Dynole 34-2 and an increase with Ryngo 1-23 (Fig 5B), supporting a role for Dnm2 during this stage of phagophore formation.

**Figure 5.**
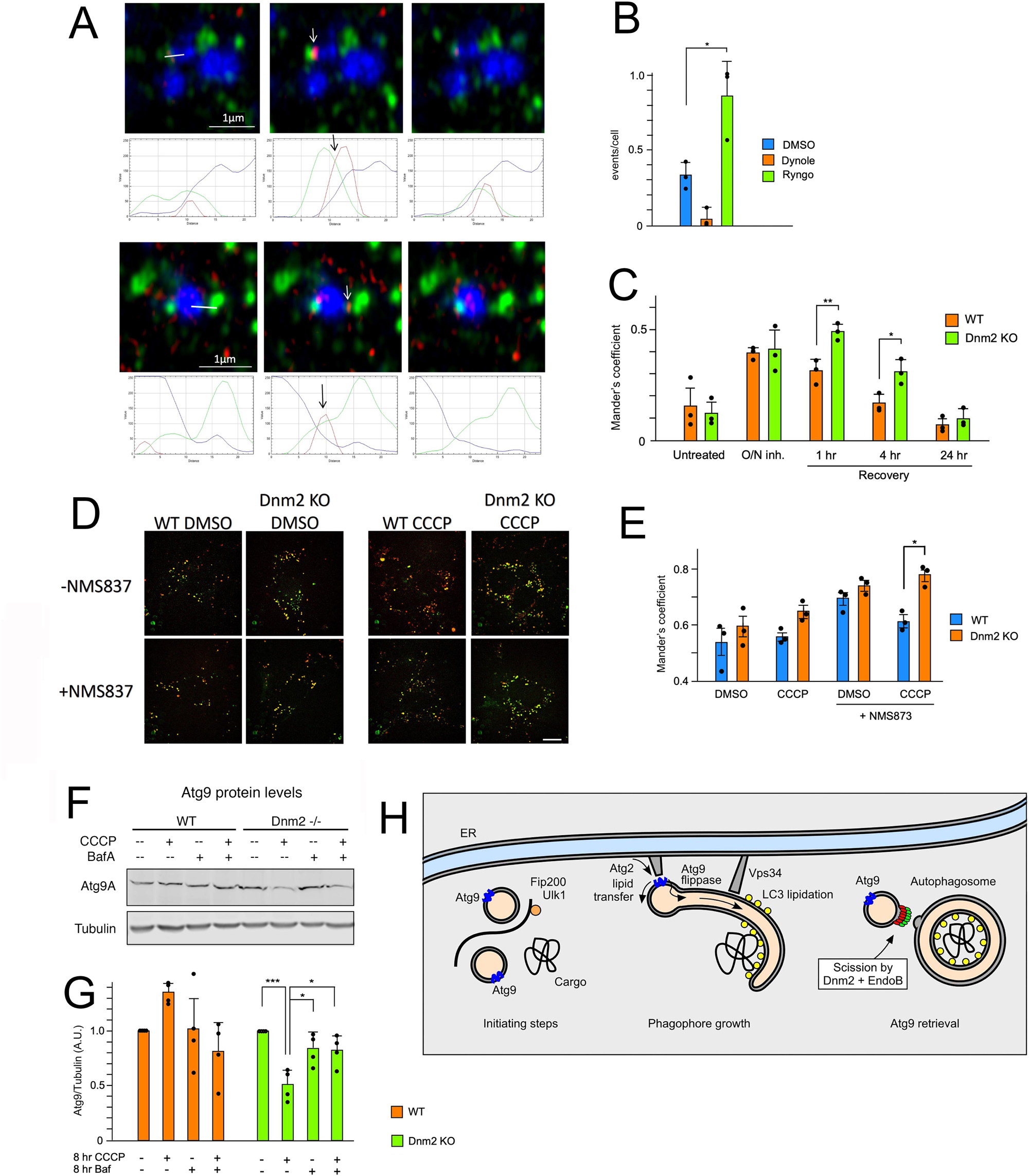
Dnm2 promotes Atg9A retrieval from phagophores. (**A**) Examples of Dnm2-RFP wedged between spots with GFP-Atg9A, and BFP-LC3 with fluorescence intensities for the transect plotted below. Frames in live cell imaging were separated by 350-400 msec. Scale bar is 1 μm. **(B)** Mean frequencies of Dnm2-RFP spots transiently wedged between GFP-Atg9A and BFP-LC3 spots in untreated cells and cells treated with Dynole 34-2 or Ryngo 1-23. The frequencies were low because only a small part of a cell was observed for a limited time (20 sec), but their transient nature and the opposing effects of a dynamin inhibitor and an activator suggest a functional connection with fission events that separate Atg9A and LC3. Average and SD are shown for three biological replicates, each determined with 4-7 technical repeats. Statistical analysis used a one-way ANOVA with Tukey’s post hoc comparisons across biological replicates. **(C)** Delayed dissociation of Atg9A and p62 in Dnm2 KO cells. A buildup of Atg9A and p62 was induced for 16h with the Vps34 inhibitor Vps34-IN1. This was followed by inhibitor washout for the indicated durations. Association and dissociation of Atg9A and p62 were monitored by immunofluorescence microscopy of endogenous proteins in WT and Dnm2 KO HeLa cells. These cells were fixed with paraformaldehyde before imaging. Average and SD are shown for three biological replicates, each determined with five technical repeats. Statistical analysis used an unpaired Student’s t-test on the biological replicates for each treatment. **(D,E)** Colocalization of RFP-Atg9A and GFP-LAMP1 after treatment with CCCP is increased by adding the p97 inhibitor NMS-873. Cells were incubated for 4 hr with DMSO or 10 μM NMS-973 and DMSO or 10 μM CCCP as indicated. Scale bar is 10 μm. Average and SD are shown for three biological replicates, each determined with five technical repeats. Statistical analysis used one-way ANOVA with Tukey’s post hoc comparisons of the biological replicates for each treatment. **(F,G)** Western blot showing decreased amounts of Atg9A protein in CCCP-treated Dnm2 KO cells. The lower panel shows quantification of Atg9A protein levels in blots of wild-type and Dnm2 KO cells. Levels decrease in Dnm2 KO cells after treatment with CCCP, but this decrease is partially prevented by Bafilomycin A, suggesting that Atg9A is degraded by autophagy in Dnm2 KO cells. Band intensities were determined with Licor software. Average and SD are shown for four biological replicates. Statistical analysis used one-way ANOVA with Tukey’s post hoc comparisons of the biological replicates for each cell line. **(H)** Functions of Dnm2 and EndoB1 on phagophores. Autophagy is triggered by activation of ULK1 kinase, which is associated with the scaffold protein FIP200 and with Atg9A-containing vesicles. The phagophore then grows from the Atg9A vesicles by Atg2-mediated lipid transfer from other organelles, such as the ER, and these lipids are equilibrated between the leaflets of the phagophore membrane by the scramblase function of Atg9A. Vps34-mediated lipid phosphorylation and subsequent LC3 lipidation promote phagophore encapsulation of cargo. Once the phagophore is complete, EndoB1 and Dnm2 mediate fission to form a vesicle that retrieves Atg9A. For the sake of simplicity, only a subset of autophagy proteins is shown.

We used a pulse-chase approach to study the timing of Atg9A trafficking in the absence of Dnm2. LC3 lipidation was blocked for 16 hours using a Vps34 inhibitor (Vps34-IN1), then washout and recovery periods of increasing lengths followed. We tracked Atg9A and p62 colocalization with immunofluorescence in WT and Dnm2 KO HeLa cells, noting that Atg9A immunofluorescence was more specific in human cells than in mouse cells. Both WT and Dnm2 KO cells showed increased colocalization of Atg9A and p62 after 16 hours of Vps34-IN1 treatment (Fig 5C), indicating clustering of Atg9A vesicles near stalled phagophores. This colocalization gradually decreased over 24 hours after removing Vps34-IN1, but recovery was notably slower in Dnm2 KO cells (Fig 5C), suggesting impaired Atg9A retrieval.

We also examined the colocalization of Atg9A and Lamp1, a lysosomal protein usually not associated with Atg9A vesicles. Cells were treated for 4 hr with or without 10 μM of the p97 inhibitor NMS-873 to prevent the extraction of Atg9A from lysosomes and degradation by the proteasome, along with 10 μM CCCP to induce mitophagy. The results showed a significant increase in colocalization of Atg9A and LAMP1 in Dnm2 KO cells following CCCP induction (Figs 5D-E). Similarly, enhanced colocalization was observed with pepstatin and E64d to prevent lysosomal degradation of Atg9A (Fig S4F-G), suggesting that Atg9A on amphisomes can be internalized and then degraded in autophagolysosomes by internal proteases as well. The extraction of ubiquitinated Atg9A by p97 would act in parallel with this degradative pathway.

We conclude that a greater proportion of Atg9A is degraded in DNM2 KO cells treated with CCCP than in WT cells.

Lastly, we used Western blots to monitor Atg9A levels in Dnm2 KO cells. The Atg9A protein levels decrease when mitophagy is induced in Dnm2 KO cells with CCCP, but not in wild-type cells (Fig 5F-G). This decrease is halted with Bafilomycin, suggesting that Atg9A is degraded by lysosomal degradation in Dnm2 KO cells, but not in wild-type cells. Together, these experiments demonstrate that Atg9A retrieval from phagophores is impaired in Dnm2 KO cells and that Atg9A is subsequently degraded in lysosomes. Although the overall rates of autophagy do not decrease in Dnm2 KO cells, there may be other functional consequences because some early autophagy proteins, such as Atg9A and SNAREs, may be turned over more rapidly than in wild-type cells, and their retention on phagophores could interfere with downstream processes. Collectively, our results suggest that Dnm2 and EndoB mediate membrane fission to retrieve Atg9A from phagophores before autophagosomes are completed and fuse with lysosomes (Fig 5H).

## Discussion

Our data reveal an unexpected role for Dnm2 in autophagy. We find that Dnm2 facilitates the retrieval of Atg9A from growing phagophores, consistent with the previously reported absence of Atg9A from mature autophagosomes (Orsi A et al, 2012). Our findings provide five lines of evidence demonstrating that Dnm2 is essential for this retrieval process. First, Dnm2 and EndoB colocalize with each other and with autophagy markers like LC3 during autophagy induction. Second, mitophagy and starvation-induced macroautophagy proceed normally in cells lacking Dnm2 and EndoB. Third, we observe significant colocalization of Atg9A with LC3 in Dnm2 KO cells and in extracts upon inducing autophagy. Additionally, SNARE proteins involved in the fusion of Atg9A vesicles with phagophores also merge with LC3 under these conditions in Dnm2 knockout cells. Similar effects are seen using other methods to impair Dnm2, such as siRNA knockdown. These results suggest that transmembrane proteins in Atg9A vesicles colocalize with LC3-containing membranes when Dnm2 function is disrupted. Fourth, Dnm2 appears in transient spots between Atg9A- and LC3-positive compartments, and chemical inhibitors or activators of Dnm2 modulate the number of these spots in live cells. Fifth, the number of Atg9A vesicles decreases when autophagy is induced in Dnm2 knockout cells, indicating that Atg9A vesicles fail to reform after seeding phagophores. Moreover, Atg9A protein levels are reduced in Dnm2 KO cells and decrease further upon induction of autophagy. This reduction is prevented by Bafilomycin, suggesting that Atg9A is incorporated into autophagosomes and degraded via autophagy when Dnm2 function is impaired. Overall, these findings show that Dnm2 is necessary for recycling transmembrane proteins from phagophores.

The new role of Dnm2 and EndoB in retrieving Atg9A from phagophores appears to conflict with previous ideas where Dnm2 separates Atg9A-containing vesicles from recycling endosomes, either with EndoB1 (Takahashi Y et al, 2016) or with SNX-18 (Soreng K et al, 2018) during starvation. In those cases, mutations in Dnm2 and EndoB1 would block Atg9A from reaching the sites of phagophore formation, leading it to accumulate in other compartments, which contradicts our findings. Different origins of phagophores could explain some of these differences. For instance, it has been shown that phagophores can grow from recycling endosomes (Puri C et al, 2018), while earlier research suggested they originate from or near the ER (Karanasios E et al, 2016, Orsi A et al, 2012). The ER proximity explanation aligns with the newly discovered roles of Atg2 and Atg9A in expanding phagophores from Atg9A vesicles close to the ER or other organelles (Chang C et al, 2021, Noda NN, 2021), and it supports a role for Dnm2 in retrieving Atg9A from those phagophores.

Dnm2 and EndoB1 may also act as barriers to prevent Atg9A from dispersing along expanding phagophores, while still allowing lipids to flow freely from transfer sites with Atg2 and Atg9A to the rest of the phagophore. Interestingly, this function appears to be absent in yeast, where Atg9A is transported to the vacuole and then retrieved (Yamamoto H et al, 2012). Other aspects of the fission process have not yet been characterized, but some similarities can be observed with Dnm2-driven fission events at the plasma membrane. In addition to Dnm2’s well-known roles in clathrin- and caveolin-dependent endocytosis (Henley JR et al, 1998, van der Bliek AM et al, 1993) and clathrin/caveolin-independent endocytosis (Conner SD & Schmid SL, 2003), Dnm2 also facilitates fast endocytosis alongside Endophilin A (FEME) (Renard HF et al, 2015). Whether EndoB1 and Dnm2 have roles in autophagy similar to Dnm2 and Endophilin A in FEME remains unclear. However, the effects of the dynamin activator Ryngo 1-23 and the inhibitor Dynole 34-2 on the appearance of Dnm2 spots between Atg9A and LC3 membranes (Fig 5B), as well as the impact of Dnm2 loss of function on Atg9A retrieval, support Dnm2’s involvement in scission at autophagosomes (Fig 5H).

In conclusion, our data suggest a model where Dnm2 facilitates Atg9A retrieval from expanding phagophores. Other possible roles in related processes, such as vesicle budding from endosomes (Soreng K et al, 2018, Takahashi Y et al, 2016) or retrieval from autolysosomes (Fang X et al, 2016, Schulze RJ et al, 2013), as well as indirect effects from different vesicular transport pathways, are also plausible.

However, our findings are most straightforwardly interpreted as direct effects on the retrieval of Atg9A-containing vesicles.

## Materials and methods

### Plasmids

The pMito-DsRed2 plasmid was from Clontech. Addgene provided pcDNA3-mRuby2 (#40260), GFP-C1-PLCdelta-PH (#21179), Dnm2-EGFP (#34686), Dnm2-mCherryN1 (#27689), ptfLC3 (#21074), pMXs-puro-RFP-ATG9AA (#60609), pEGFP-LC3 (# 24920), GFP-Rab7A (#61803), mTaqBFP2-ER-5 (#55294), pcDNA3-mRuby2 (#40260), pEGFP-C1-hAtg2A (#36456), pMXs-IP-EGFP-hFIP200 (#38192), pEGFP-VAMP7 (#42316), pmRFP-LC3 (#21075) and pMRXIP GFP-Stx17 (#45909). X. Soriano (Department of Cell Biology, University of Barcelona) provided the GFP-Bax plasmid. E. C. Dell’Angelica (Department of Human Genetics, UCLA School of Medicine) provided the GFP-Lamp1 plasmid. To generate EndophilinB1- mRuby2, EndophilinB1 coding sequences from cDNA were fused in frame to mRuby2 by PCR cloning into pcDNA3-mRuby2. EGFP-Atg9A, Dnm2-RFP and mTaqBFP2-LC3 constructs were generated in the same way with coding sequences from pMXs-puro-RFP-ATG9AA, Dnm2-EGFP and mTaqBFP2-ER-5, respectively.

### Cell culture, transfection, gene knockout and chemical treatments

WT HeLa cells were from James Wohlschlegel (Dept. of Biological Chemistry, UCLA), WT MEFs were from David Chan (Dept of Biology, CalTech) and MEFs with stable expression of FLAG-Parkin were from Lars Dreier (UCLA) (Sun Y et al, 2012). All cell lines were periodically checked for Mycoplasm. MEFs were grown in DMEM with 10% FBS. Transient transfections were done with jetPRIME following manufacturer’s instructions (Polyplus). For siRNA, cells were grown in 6cm dishes, transfected with 50nM oligonucleotides using RNAimax (Invitrogen) and analyzed 72h later. Deletions in the Dnm2 and EndoB1 gene were introduced in MEFs with FLAG-Parkin using CRISPR/Cas9 methods, as described (Ran FA et al, 2013). Target sites were identified using the Boutros Lab Website (http://www.e-crisp.org/E-CRISP/designcrispr.html). Dnm2 gRNA was: 5’-GATGGCAAACACGTGCTTGT-3’. EndoB1 gRNAs were: 5’-GCAGGAACTGAGTTTGGCCC-3’; and 5’-GGATTTCAACGTGAAGAAGC-3’. The gRNAs were cloned in the px459 plasmid, containing puromycin resistance and Cas9 genes, and 1.2μg was transfected into MEF cells, followed by 24hr puromycin selection. Surviving colonies were isolated, genotyped and analyzed with western blots. HeLa cells were also grown in DMEM with 10% FBS. The gRNA used to knock out Dnm2 in HeLa cells was: 5’-ACAGGGGCCGGCCCTGGACC-3’. The knockout procedure was as described for MEF cells.

Because the EndoB1 KO cells lacked Parkin expression, they were stably transfected with an EF1alpha promoter-driven Parkin expression construct in a PiggyBac vector. This construct was transfected into EndoB KO MEFs along with the PBase vector from Vectorbuilder at equal concentrations (2µg/per well in a 6 well plate). Atg9A-myc and HA-LC3B constructs were similarly stably transfected using the PiggyBac protocol. At 24h post-transfection, cells were subjected to puromycin selection at increasing concentrations from 1µg to 5 µg/ml for 1-2 days. After selection, surviving colonies were isolated with cloning rings, expanded and analyzed for Parkin expression levels by Western blotting. A clone with Parkin expression levels that matched those in WT with FLAG-Parkin cells and Dnm2 with FLAG-Parkin cells was selected for further analysis.

Amino acid starvation was induced by washing cells twice in EBSS with calcium and magnesium (Thermo Fisher Scientific) and incubating them for four hours. The following chemicals with their final concentrations were from Sigma-Aldrich: 20μM CCCP, 40μg/ml antimycin A, 10μM actinomycin D and 1μM 4-Hydroxy Tamoxifen. The following chemicals with their final concentrations were from InvivoGen: 0.5μM Bafilomycin A1, and 10μM rapamycin. Staurosporine (Thermo Fisher Scientific) was used at 2μM. Z-VAD-FMK (BD Bioscience) was used at 20μM. Dynasore (EMD Millipore) was used at 80μM. VPS34-IN1 (Selleckchem) was used at 10μM. NMS-873 (Sigma-Aldrich) was used at 10μM. Pepstatin and E64d (Sigma-Aldrich) were used at 5 µg/ml.

### Immunoblotting, immunofluorescence, and proximity ligation assay

Total cell lysates for Western blots were made with RIPA buffer. Samples were subjected to SDS-PAGE, transferred to nitrocellulose or PVDF membranes, blocked with Odyssey Blocking Buffer (LI-COR), and incubated overnight at 4°C with primary antibodies. Membranes were then washed with PBS-T and incubated with IRDye 800CW or 670RD secondary antibodies (LI-COR). Fluorescent bands were detected with an Odyssey scanner and analyzed with Image Studio Software (LI-COR). For immunofluorescence images, cells were grown on 12mm coverslips, fixed for 10min with 4% paraformaldehyde in PBS, and permeabilized for 5min with 0.1% Triton X-100 in PBS, blocked for 1h with Goat or Donkey Serum in PBS-T and incubated with primary antibodies. One exception was immunofluorescence of Dnm2 (Figs 1A,B), for which cells were first permeabilized with 0.05% digitonin to wash out soluble cytosolic proteins (Liu SH et al, 1998), before fixation with 4% paraformaldehyde. Secondary antibodies were Alexa Fluor 488-, 594- or 647-conjugated goat anti-mouse or rabbit IgG (Invitrogen). Proximity ligation assays were conducted with Duolink as recommended by the manufacturer (Sigma-Aldrich). Cells expressing fluorescent proteins were also fixed with 4% paraformaldehyde, except when live cells were imaged, as indicated in the figure legends.

### Antibodies

Rabbit anti-Endophilin A1, mouse anti-SQSTM1 (p62), rabbit anti-dynamin 1, rabbit anti-dynamin 2, mouse anti-Hsp60, rabbit anti-cytochrome c and mouse anti-VDAC1/Porin were from Abcam. Rabbit anti-LC3B and mouse anti-tubulin antibodies were from Sigma Aldrich. Goat anti-dynamin 2 and rabbit anti-Tom20 were from Santa Cruz Biotechnology. Mouse anti-Tim23 and mouse anti-Drp1 were from BD Biosciences. Rabbit anti-RAB7A was from Proteintech. Mouse anti-Endophilin B1 was from Imgenex. Rabbit anti-SNX18 and rabbit anti-ATG9AA were from GeneTex.

### Microscopy

Standard immunofluorescence and live cell images were made with a Marianas spinning disc confocal from Intelligent Imaging, which uses an Axiovert microscope (Carl Zeiss Microscopy) with 40x/1.4 and 100x/1.4 oil objectives, a CSU22 spinning disk (Yokogawa), an Evolve 512 EMCCD camera (Photometrics) and a temperature unit (Okolab). Where it is indicated that live cells were imaged, the cells were grown in glass-bottom dishes (MatTek) and imaged in situ at 37°C. Super-resolution images were acquired with SIM using a DeltaVision OMX SR (General Electric). Fiji ImageJ software was used for image analysis. Time lapse images of live cells were acquired with a Zeiss Airyscan microscope equipped with a Fast-Airyscan module and processed with Airyscan software.

### Subcellular fractionation

Extracts for imaging of partially purified autophagosomes were essentially made as described (Gao W et al, 2010). In brief, MEF cells were transfected with EGFP-Atg9A and RFP-LC3. After 16h, these cells were treated for 4 h with 20 μM CCCP and 0.5 μM Bafilomycin. Cells were harvested in mitochondrial isolation buffer (0.25 M sucrose, 1 mM EDTA, 20mM HEPES, pH 7.4) on ice, disrupted by passing 20 times through a 22-gauge needle. Nuclei and unbroken cells were removed with low-speed centrifugation (10 min at 600g) followed by enrichment of autophagosomes with medium-speed centrifugation (20 min at 10,000g) and washing twice in 1.0 M NaCl in mitochondrial isolation buffer with protease inhibitor to dissociate clusters of vesicles that were potentially held together by ionic interactions. Samples of the resulting fractions were mounted on slides and examined by fluorescence microscopy.

For Ficoll gradients, WT and Dnm2 KO MEF cells were treated for 4 hr with 0.5mM Bafilomycin A, followed by 2 hr with 0.5mM Bafilomycin A and 5 μM Rapamycin to induce autophagy. Cells were then fractionated as described (Orsi A et al, 2012), but with a few modifications. In brief, cells from 4 confluent 15 cm dishes were washed, scraped, and resuspended in homogenization buffer (250 mM sucrose, 20 mM HEPES/KOH pH 7.4, 0.5 mM PMSF), followed by homogenization with 15 passes through a 25 G needle.

Nuclei and unbroken cells were removed with low-speed centrifugation. The equivalent of 2 mg protein was loaded on a 1-22% Ficoll gradient with a total volume of 10 ml layered on a 1 ml 45% Nycodenz cushion in homogenization buffer. These preparations were centrifuged in a Beckmann SW41 rotor for 14.5 hr at 20 krpm at 4°C. Twenty fractions of 0.5 ml were collected, and even-numbered fractions were analyzed with Western blots of 40 μl samples.

## Acknowledgements

We are grateful for many helpful discussions and sharing of reagents with Lars Dreier, Yu (Sammy) Sun at UCLA. CMK was supported by NIH grants R01GM61721, R01GM073981 and R01DK101780. AMvdB was supported by NIH grants U01GM109764 and R01NS120690.

## Author Contributions

AC and AMR conducted experiments and helped design them. CIM, SI, JS and BR conducted experiments. CMK was a co-supervisor and contributed to the design of the experiments. AMvdB supervised the project, assisted in designing the experiments, and wrote the paper.

---

**Suppl. Figure 1.**
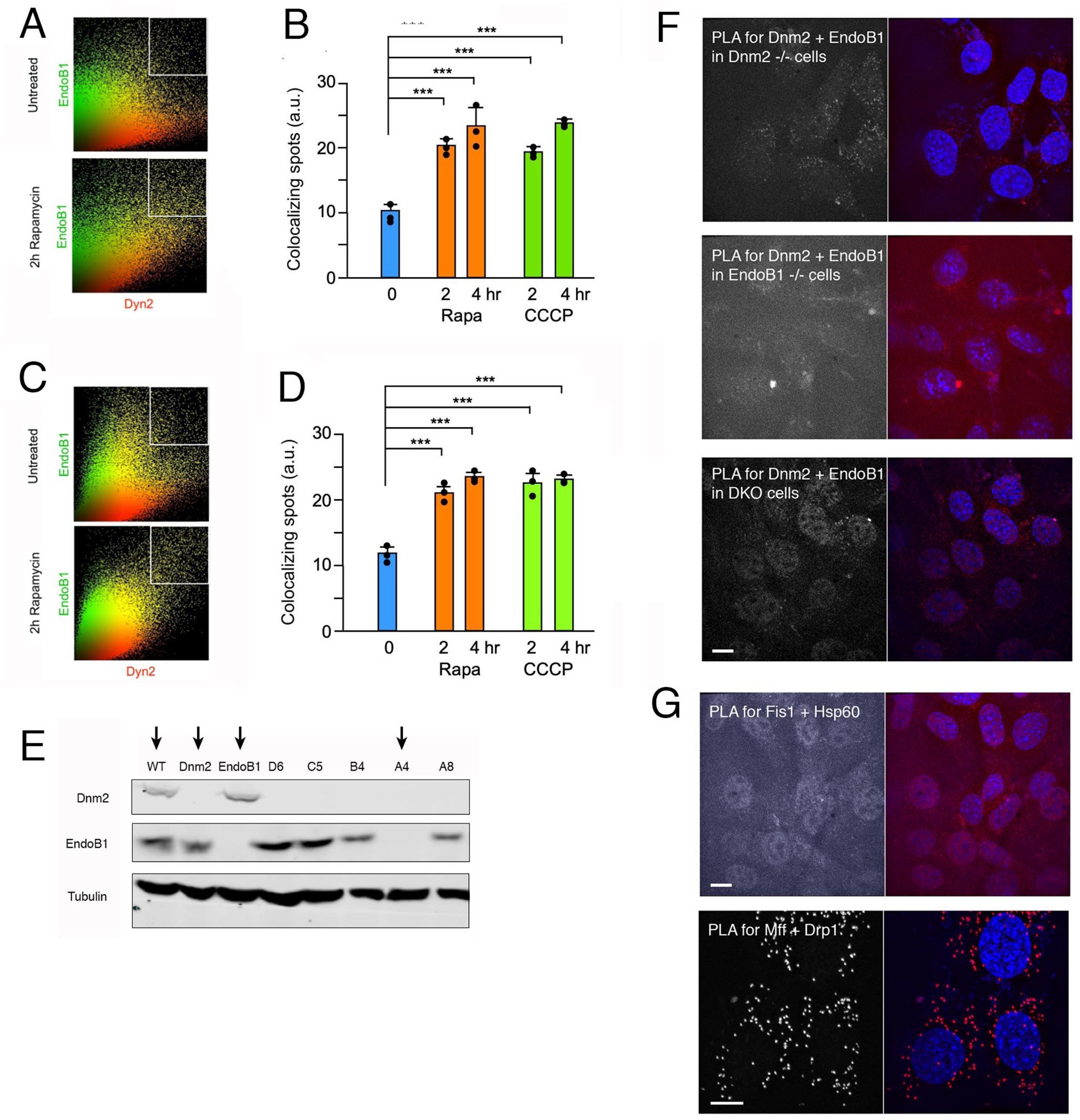
Colocalization of Dnm2 and EndoB1 shown with scatterplots. **(A)** Scatterplots of immunofluorescence images of cells treated for 2h with or without rapamycin to induce autophagy. **(B)** Fraction of spots in the colocalizing quadrants of the scatterplots, shown before and after 2 or 4 hr treatment with rapamycin or CCCP. Average and SD are shown for three biological replicates, each determined with five technical repeats. Statistical analysis used one-way ANOVA with Tukey’s post hoc comparisons of the biological replicates. **(C,D)** As in panels A and B, but with transiently expressed GFP-and RFP-tagged proteins. **(E)** Western blots showing the derivation of Dnm2, EndoB1 and DKO cells (clone A4). Dnm2 and EndoB1 KO cells were generated by CRISPR/Cas9 in MEFs. DKO cells were generated by additional knockout of EndoB1 in Dnm2 KO cells and clone A4 was chosen for further studies. **(F)** PLA signals for Dnm2 and EndoB1 interactions were absent from cells with mutations in one or both of these proteins. Scale bar is 10 μm. **(G)** As a negative control for PLA, cells were incubated with Fis1 and Hsp60 antibodies, which detect proteins that are on the outside or inside of mitochondria and are therefore too far apart for generating PLA signals. As a positive control, cells were incubated with antibodies for Drp1 and Mff, which detect well-characterized binding partners. Scale bar is 10 μm.

**Suppl. Figure 2.**
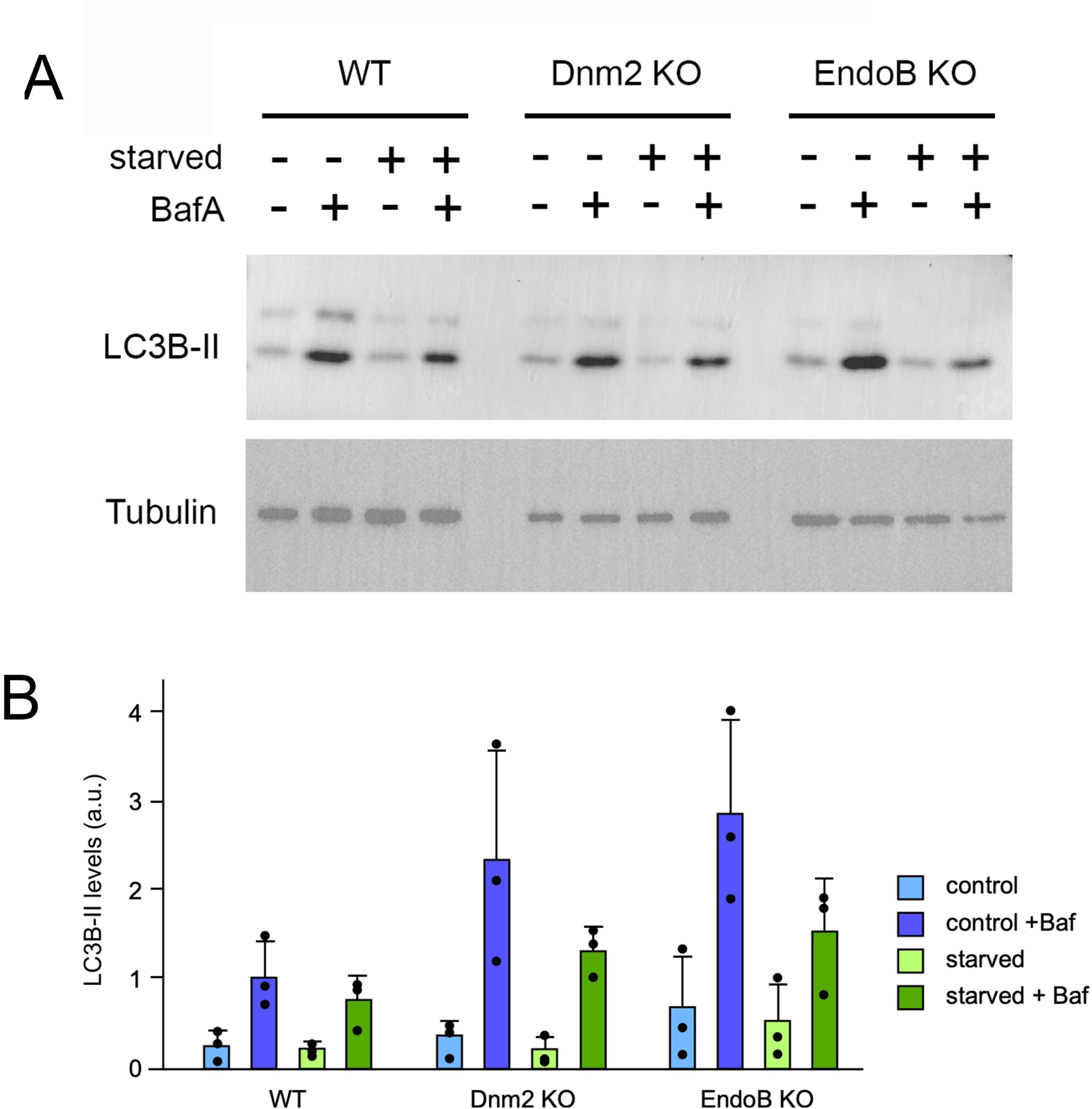
Effects of EndoB KO on mitophagy in MEFs and of Dnm2 and EndoB KO on starvation-induced LC3B lipidation. **(A)** Western blots showing the effects of starvation-induced macroautophagy and Bafilomycin A treatments on LC3B lipidation in wildtype, Dnm2 and EndoB KO HeLa cells. **(B)** Average intensities of LC3B-II bands in four independent experiments, relative to tubulin levels, expressed in arbitrary units (a.u.). Average and SD are shown for three biological replicates. Statistical analysis using one-way ANOVA with Tukey’s post hoc comparisons did not show significance.

**Suppl. Figure 3.**
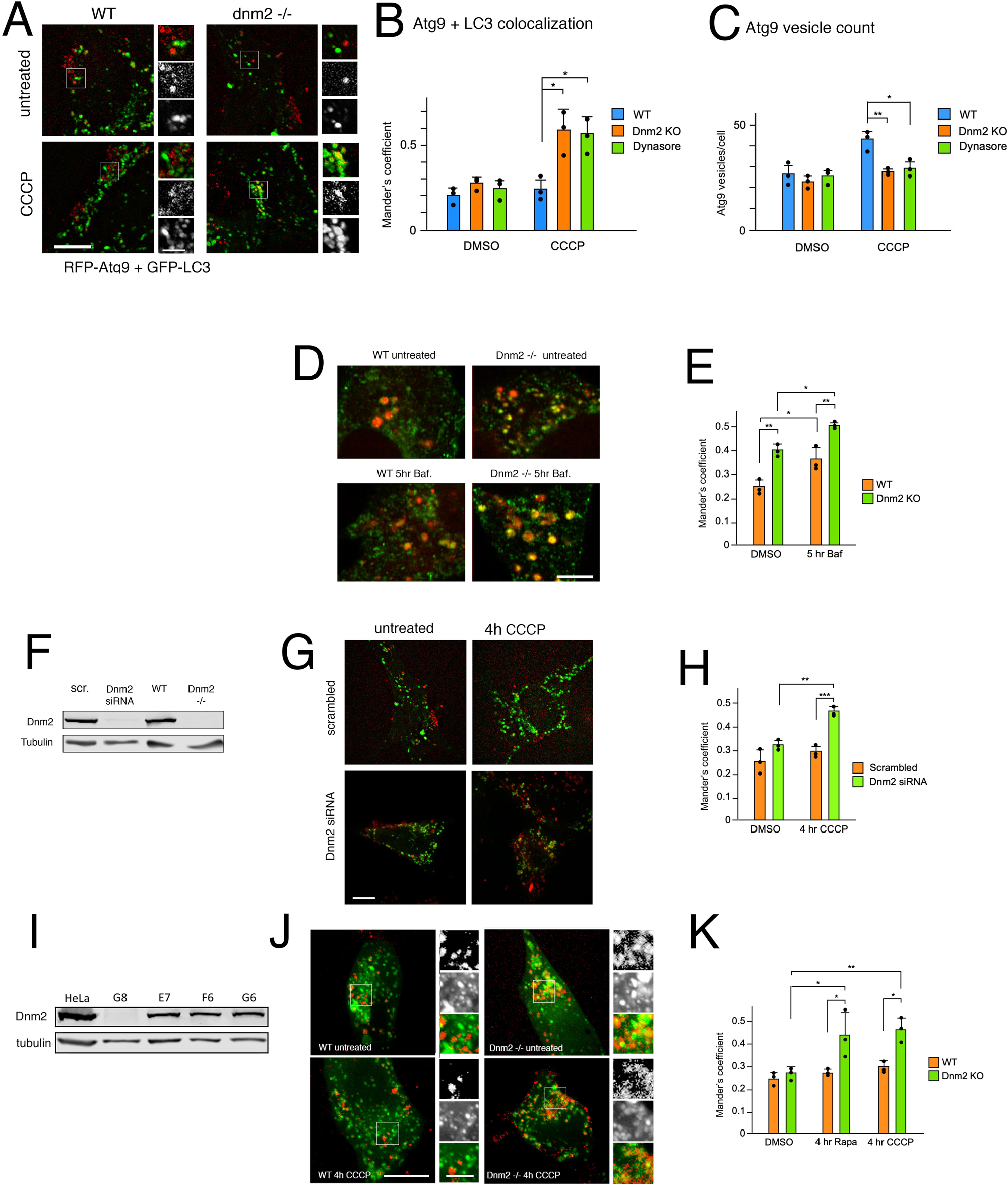
Atg9A localization in wt and Dnm2 siRNA MEF cells and in Dnm2 KO HeLa cells. **(A)** Live cell fluorescence images show RFP-Atg9A colocalizes with EGFP-LC3 in Dnm2 KO MEFs treated with CCCP, but not in wild type MEFs. Scale bar is 10 μm for whole cells and 3 μm for enlarged portions. **(B)** Histogram showing Mander’s coefficient for colocalization of RFP-ATG9A and GFP-LC3 in Dnm2 KO MEFs and dynasore-treated WT MEFs after incubation with rapamycin or CCCP. Average and SD are shown for three biological replicates, with five technical replicates per biological replicate. Statistical analysis used one-way ANOVA with Tukey’s post hoc comparisons. **(C)** The number of Atg9A vesicles (RFP spots without GFP label) decreases when Dnm2 KO MEFs and dynasore-treated WT MEFs are incubated with rapamycin or CCCP. Statistics as in panel B. **(D,E)** Effects of Dnm2 on Atg9A retrieval in live cells. Images of BFP- LC3 (shown in red) and GFP-Atg9A in live WT and Dnm2 KO cells, untreated or after 5 hr with Bafilomycin. Scale bar is 5 μm. Histograms showing increased colocalization of BFP-LC3 and GFP-Atg9A in Dnm2 KO cells with or without Bafilomycin treatments. Average and SD are shown for three biological replicates with five technical replicates for each one. Statistical analysis used one-way ANOVA with Tukey’s post hoc comparisons. Scale bar is 10 μm. **(F)** Western blot showing depletion of Dnm2 after transfection of WT cells with siRNA oligonucleotides. **(G)** RFP-Atg9A colocalizes with EGFP-LC3 after 4h treatment with CCCP in Dnm2 siRNA cells, but not in mock-transfected cells. Scale bar is 10 μm. **(H)** Histogram showing increased colocalization of RFP-ATG9A and GFP-LC3 in CCCP-treated Dnm2 siRNA cells. Average and SD are shown for three biological replicates with five technical replicates for each one. Statistical analysis used one-way ANOVA with Tukey’s post hoc comparisons. **(I)** Western blot showing depletion of Dnm2 in a CRISPR/Cas9 generated HeLa cell clones. Clone G8 was chosen for further analysis. **(J)** RFP-Atg9A colocalizes with EGFP-LC3 in Dnm2 KO HeLa cells treated with CCCP, but not in wild type cells. Scale bar is 10 μm for whole cells and 3 μm for enlarged portions. **(K)** Histogram showing increased colocalization of RFP-ATG9A and GFP-LC3 in Dnm2 KO HeLa cells after incubation with rapamycin or CCCP. Average and SD are shown for three biological replicates, with five technical replicates per biological replicate. Statistical analysis used one-way ANOVA with Tukey’s post hoc comparisons.

**Suppl. Figure 4.**
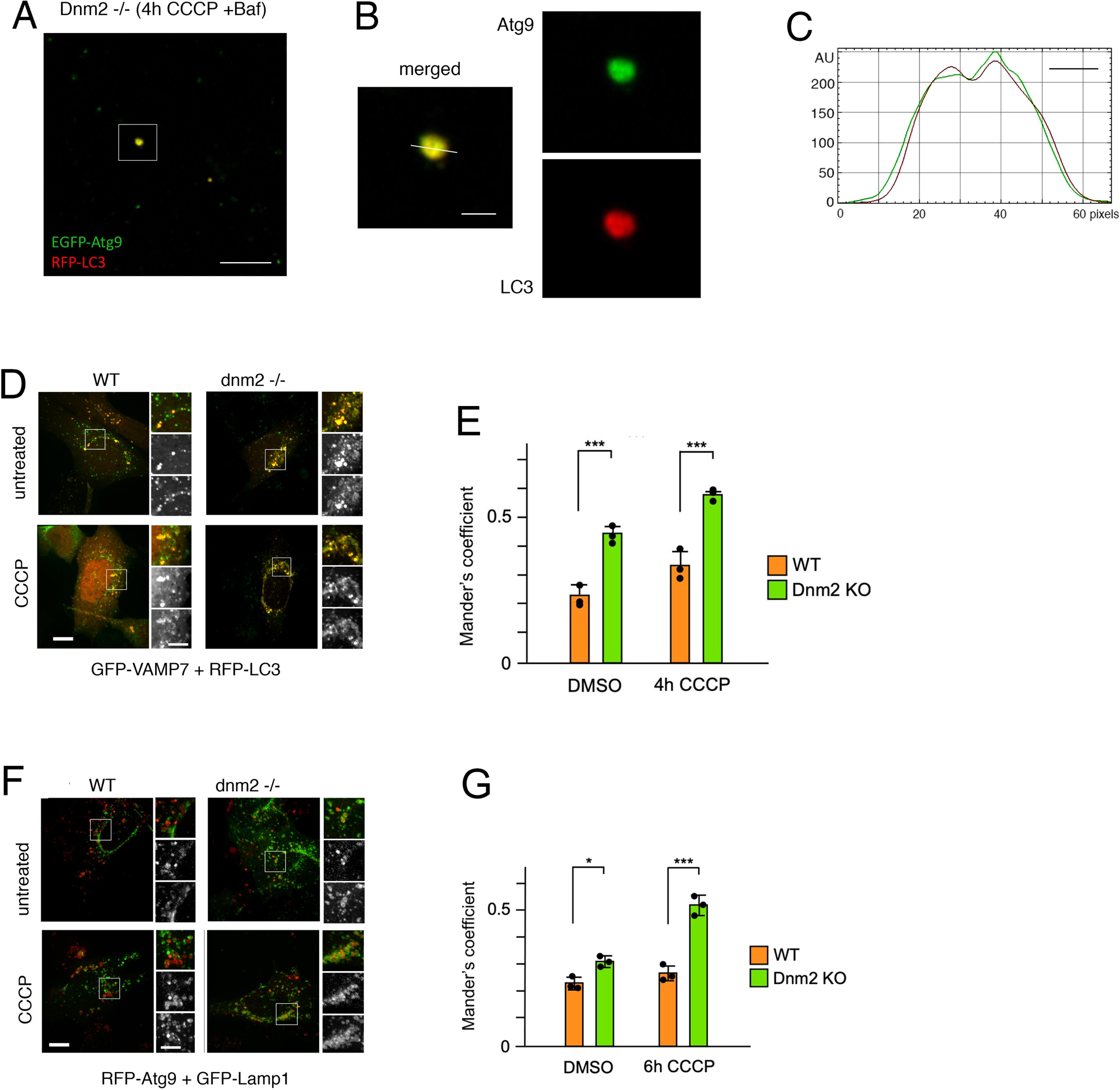
Colocalization of GFP-Atg9A and RFP-LC3 in autophagosome fractions and of GFP-VAMP7 and RFP-LC3 in live cells. **(A)** Image of autophagosome-containing fraction from Dnm2 KO cells transfected with EGFP-Atg9A and RFP-LC3, followed by 4hr treatments with CCCP and Bafilomycin to induce mitophagy and to accumulate autophagosomes. Autophagosomes were enriched by differential centrifugation. Scale bar is 10 μm. **(B, C)** Enlargements show an individual spot with the line used to plot intensities. Scale bar is 2 μm. The x-axis in the fluorescence intensity plot shows numbers of pixels (24 pixel/μm) and the bar shows 500 nm. **(D, E)** Colocalization of GFP-VAMP7 with RFP-LC3 is higher in Dnm2 KO MEFs than wildtype cells upon treatment with CCCP. Scale bar is 10 μm. Average and SD are shown for three biological replicates, with five technical replicates per biological replicate. Statistical analysis used one-way ANOVA with Tukey’s post hoc comparisons. (**F, G**) There is also increased colocalization between RFP-Atg9 and GFP-LAMP1 after CCCP treatment. Cells were incubated with pepstatin and E64d to prevent Atg9A digestion in lysosomes. Scale bar is 10 μm for whole cells and 5 μm for enlarged portions. Average and SD are shown for three biological replicates, with five technical replicates per biological replicate. Statistical analysis used one-way ANOVA with Tukey’s post hoc comparisons.

